# A mixture of three engineered phosphotriesterases enables rapid detoxification of the entire spectrum of known threat nerve agents

**DOI:** 10.1101/748749

**Authors:** Dragana Despotovic, Einav Aharon, Artem Dubovetskyi, Haim Leader, Yacov Ashani, Dan S. Tawfik

## Abstract

Nerve agents are organophosphates that potently inhibit acetylcholinesterase and their enzymatic detoxification has been a long-standing goal. Nerve agents vary widely in size, charge, hydrophobicity, and the cleavable ester bond. A single enzyme is therefore unlikely to efficiently hydrolyze all agents. Here, we describe a mixture of three previously developed variants of the bacterial phosphotriesterase (*Bd*-PTE) that are highly stable and nearly sequence identical. This mixture enables effective detoxification of a broad spectrum of known threat agents – GA (tabun), GB (sarin), GD (soman), GF (cyclosarin), VX, and Russian-VX. The potential for dimer dissociation and exchange that could inactivate *Bd*-PTE has minimal impact, and the three enzyme variants are as active in a mixture as they are individually. To our knowledge, this engineered enzyme ‘cocktail’ comprises the first solution for enzymatic detoxification of the entire range of threat nerve agents.

## Introduction

Rapid detoxification and/or decontamination of organophosphate nerve agents (OPNAs) is highly desirable yet not readily achievable. Catalytic bioscavengers have and are being developed. Specifically, organophosphate hydrolases (OPHs) can hydrolyze and thereby inactivate OPNAs, and can do so at high rate at ambient conditions [1–3]. Enzymes, however, are generally specific, and accordingly enzyme variants that degrade one, or few related OPNAs have been reported. However, detoxification of the entire spectrum of known threat OPNAs by a single enzyme, and with high catalytic efficiency, remains an unmet challenge. The snag is that OPNAs differ widely in size, hydrophobicity and charge (Figure 1). Foremost, the two major classes of OPNAs, G- and V-agents, fundamentally differ in their leaving groups, thus making the task of complete coverage by a single enzyme even more challenging.

**Figure 1.**
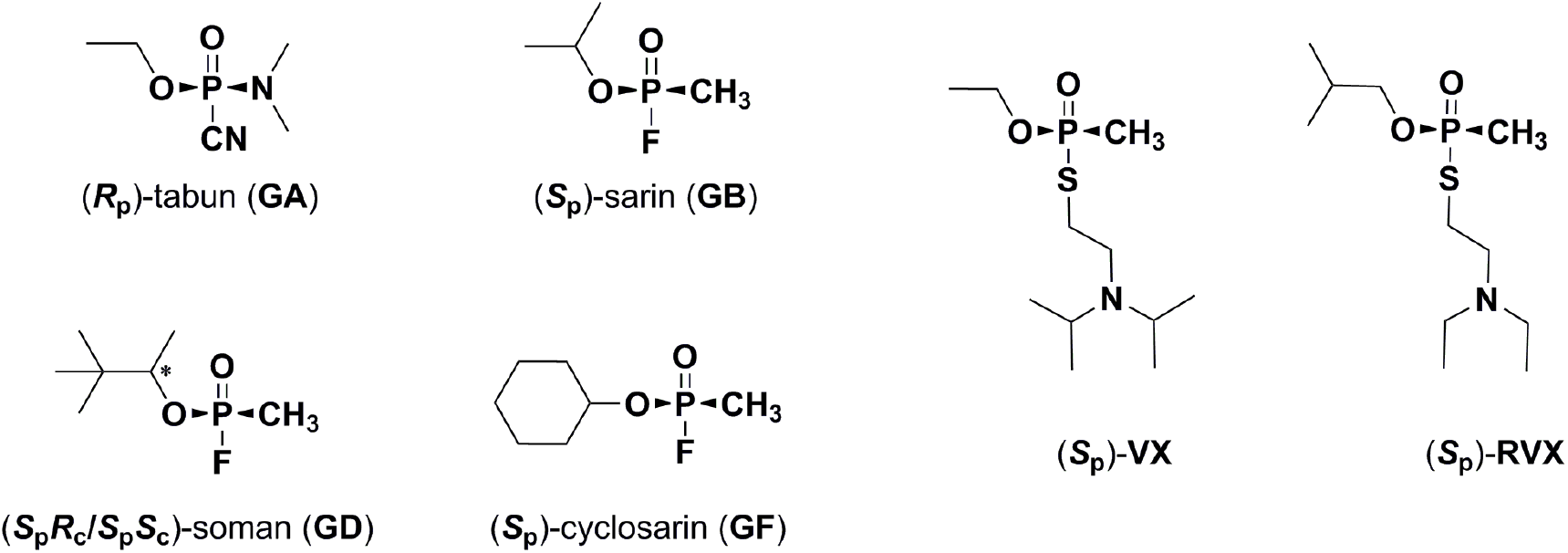
The structures of threat agents (GA, GB, GD and GF, and VX and RVX). The absolute configuration of the toxic isomers in noted in the agent names. The leaving groups released upon hydrolysis are depicted at the bottom, underneath the phosphorous (CN and F for G-agents, and thiol leaving groups for V-agents). Note that GD has an additional chiral center (the pinacolyl carbon, labeled with a star), yet both these isomers exhibit similar toxicity.

Nature has typically resolved the challenge of detoxifying a broad range of related compounds by evolving paralogs, namely several enzyme variants that have diverged from one common ancestral enzyme. Although each is specific for a certain substrate, paralogous enzymes tend to overlap in activity, thus jointly tackling a broad spectrum of toxic compounds [4–6]. Here, we have adopted this strategy for nerve agent detoxification, by combining three enzyme variants that were evolved from the same parental enzyme. Each of these variants was evolved toward a specific nerve agent, or subclass of nerve agents, yet also exhibits weaker activity with other agents. When combined, an effective detoxification of the entire spectrum of known threat nerve agents could be achieved.

The phosphotriesterase isolated from *Brevundimonas diminuta* (recently reclassified as *Sphingopyxis wildii* [7]) served as the parental enzyme (*Bd*-PTE; also known as OPD) [8–10]. Among the enzymes capable of hydrolysis of OPNAs, this enzyme has been recognized as the most promising candidate for detoxification and/or decontamination. Wild-type *Bd*-PTE hydrolyzes G-agents, including the ones with branched O-alkyl groups, sarin (GB) and soman (GD), with k_cat_/K_M_ values ranging from 0.5 to 5×10^6^ M^−1^ min^−1^ [11]. However, its activity towards VX and Russian VX (RVX) is 10 to 100-fold lower [12,13]. Overall, these catalytic efficiencies are far from sufficient for practical uses, especially because for most agents the non-toxic *R*_p_ isomer is the preferred substrate [14]. Regardless of the intended use, be it skin and/or equipment decontamination, or pre- and post-exposure detoxification *in vivo*, catalytic efficiency is a make-or-brake factor. We hypothesized and experimentally confirmed that effective protection at a reasonable *in vivo* dose (<1 mg/Kg) demands a k_cat_/K_M_ value ≥ 10^7^ M^−1^ min^−1^ for hydrolysis of the toxic OPNAs isomers [15]. Similarly, enzyme-based decontamination demands low-cost, highly stable and highly catalytically efficient enzymes [16,17].

In the last decade, combined efforts of rational design, computational design and directed evolution produced a large body of improved variants of PTE, as well as of other OPHs such as serum paraoxonase 1 (PON1) and organophosphorus acid anhydrolase (OPAA), for various OPNAs [18–22]. However, sufficiently high catalytic efficiency is achieved only for the specific OPNA or structurally similar OPNAs. Our own engineering efforts yielded up to 5000-fold enhancement of wild-type PTE’s rates of hydrolysis of the toxic isomer of VX (*S*_p_-VX) thus yielding PTE variant with k_cat_/K_M_ of 5×10^7^ M^−1^min^−1^ (PTE-d1-10-2-C3) [20] (for another VX specific PTE variant, see [19]). However, this and other improved VX hydrolyzing variants exhibited much lower activity with the related Russian VX analog, RVX (k_cat_/K_M_ for *S*_p_-RVX ≤ 0.3×10^7^ M^−1^min^−1^) and for GD (0.14×10^7^ M^−1^min^−1^) [20]. Indeed, tradeoffs between substrates are widely recorded [23]. We have accordingly observed a consistent tradeoff between VX and RVX improvements, and thus, to achieve the desired catalytic efficiency with RVX, a parallel optimization trajectory had to be pursued, yielding the *S*_p_-RVX specialized variant PTE-d1-IVA1 [20]. Similarly, by screening computationally designed PTE variants, we have recently identified a third variant, *Bd*-PTE-d2-R2#16, that inactivated the toxic *S*_p_ isomers of GD and GF with k_cat_/K_M_ > 10^7^ M^−1^min^−1^, but this variant was poorly active with *S*_p_-VX and *S*_p_-RVX (k_cat_/K_M_ < 10^5^ M^−1^min^−1^) [21].

The above examples, and numerous other OPH variants described in the literature, indicate that coverage of the entire spectrum of threat OPNAs with sufficient efficacy (k_cat_/K_M_ ≥ 10^7^ M^−1^min^−1^) by a single enzyme is unrealistic. A mixture of few closely related variants of the same enzyme is, however, a more feasible option, also because such a mixture might be considered by the FDA as a single biological product [24, [https://www.fda.gov/media/125484/download]. Nonetheless, to date, a combination of sufficiently active OPH variants, that act with no mutual interference, and jointly cover the entire spectrum of threat OPNAs, has not been described. Here, we describe the *in vitro* characterization of a mixture of three nearly identical *Bd*-PTE variants that jointly neutralize all known threat OPNAs with k_cat_/K_M_ in the range of 0.2×10^7^ up to 10^8^ M^−1^min^−1^ (per each of the threat agents, and with the enzyme concentration corresponding to the total concentration of the three enzyme mixture). We further describe tag-less versions of these *Bd*-PTE variants that enable non-expensive, high-yield production of intact, stable bioscavengers.

## Material and Methods

*Bd*-PTE variants N-terminally fused to MBP were expressed and purified as previously described [20].

### Cloning, expression and purification of tag-less PTE variants

The gene encoding the previously reported PTE_dC23 variant was amplified from its pMAL vector [20], and re-cloned into pET21a vector using the NdeI and XhoI restriction enzymes. Variants d1-IVA1, d1-10-2-C3 and d2-R2#16 were amplified from the originally reported pMAL vectors and transferred to pET21a vector by MEGAWHOP PCR [25] using the above pET21a_*Bd*-PTE-dC23 plasmid as template. The pET21a plasmids were transformed to *E. coli* BL21 DE3 cells for protein expression. The gene encoding the designed d1-R2#16 variant was ordered from Twist cloned in pET29b.

Colonies of *E. coli* cells with the transformed PTE variants were used to inoculate starter cultures (LB plus 100 µg/ml Amp for pET21a constructs, and 50 µg/ml Kan for the pET29b construct) that were grown overnight (ON) at 37 °C. These cultures (1.25 ml; 1:100 dilution) were used to inoculate expression cultures of 125 ml LB medium plus 100 µg/ml Amp and 0.2 mM ZnCl_2_. The cultures were grown at 37°C to OD_600_ 0.8, expression was induced by 0.4 mM IPTG, and the cultures were further incubated ON with shaking at 20°C. Cells were pelleted (2500 x g for 30 min at 4°C), the pellet was frozen at −20 °C and resuspended in 20 ml of lysis buffer (20 mM Tris, pH 7.6, 0.1 mM ZnCl_2_, supplemented with Protease Inhibitor Cocktail (EDTA-free; ABCAM, diluted 1:1000), Benzonase nuclease (MERCK, 100 units), and lysozyme (Glentham Life Sciences Ltd, 0.4 mg/ml)). Resuspended cells were incubated for 1 h at 37 °C and lysis was completed by sonication. Cell debris was removed by centrifugation (45 min at 7500 x g at 4°C) and the supernatant was filtered through a 0.22 µm syringe filter.

The enzyme variants were subject to a 3-step purification. *In the 1*^*st*^ *step*, the filtrate was loaded on an anion exchange HiTrap Q HP 5 ml column (GE healthcare life sciences, Boston, Massachusetts), at a flow rate of 1 ml/min, using an FPLC AKTA-prime plus device (GE, Boston, Massachusetts) pre-equilibrated with buffer A_Q_ (20 mM Tris, pH 7.6, 0.1 mM ZnCl_2_). The enzyme variants were eluted by a gradient of buffer B_Q_ supplemented with 1 M NaCl (0-100 % gradient within 20 column volumes, at flow rate 5 ml/min). The paraoxonase activity of the collected fractions (samples were first diluted 10^4^-fold) was measured with paraoxon (at 0.18 mM) in activity buffer (100 mM NaCl, 50 mM Tris pH 8.0). The increase in absorbance at 405 nm was followed using a plate-reading spectrophotometer (PowerWave HT, BioTek, Winooski, Vermont). The active fractions were combined and dialyzed in SnakeSkin Dialysis Tubing 10K MWCO (Thermo Fisher Scientific, Waltham, Massachusetts) against buffer A_SP_ (100 mM ammonium acetate, 0.1 mM ZnCl_2_, pH 6.0).

*In the 2*^*nd*^ *step*, the dialyzed protein was filtered and loaded on a cation exchange HiTrap SP HP 5 ml column (GE healthcare life sciences) pre-equilibrated with 5 column volumes of buffer A_SP_ (100 mM Ammonium acetate, 0.1 mM ZnCl_2_, pH 6.0) at 1 ml/min flow. The enzyme variants were collected in the flow-through and dialyzed against buffer A_Butyl_ (500 mM ammonium sulfate, 50 mM sodium phosphate (Na_2_HPO_4_), 0.1 mM ZnCl_2_, pH 7.0).

*In the 3*^*rd*^ *step*, the dialyzed protein was filtered and loaded on a hydrophobic HiTrap Butyl HP 1 ml column (GE healthcare life sciences), pre-equilibrated with 5 column volumes of buffer A_Butyl_ (500 mM ammonium sulfate, 50 mM sodium phosphate (Na_2_HPO_4_), 0.1 mM ZnCl_2_, pH 7.0). The loading rate was 0.5 ml/min, followed by washing with 5 column volumes of buffer A_HIC_ at 0.5 ml/min. Enzyme variants were eluted by gradually increasing the percentage of buffer B_Butyl_ (10 mM Ammonium sulfate, 50 mM Na_2_HPO_4_, 0.1 mM ZnCl_2_, pH 7.0) for 10 column volumes at 0.5 ml/min. Activity in the collected fractions was measured as above. The active fractions were combined and dialyzed against storage buffer (50 mM Tris, 10 mM NaHCO_3_, 0.1 mM ZnCl_2_, 0.01% NaN_3_, pH 8.0). The purity of the purified variants was assessed by SDS-PAGE (*e.g.* Figure S1).

### Heat inactivation assay

Samples of 15 μl of 0.1 mg/ml of purified enzymes in 50 mM Tris pH 8.0, 100 mM NaCl, 0.1 mM ZnCl_2_, were incubated in a PCR gradient machine (SENSOQUEST, Göttingen, Germany) for 30 min at various temperatures, and then placed on ice. The samples were subsequently diluted 10^4^-fold in activity buffer and assayed with 0.18 mM paraoxon as above. Initial rates were derived and normalized to the rate of the sample incubated at 25 °C. For the purpose of displaying, the normalized rates were fitted to a 4-parameter Boltzmann sigmoid using the GraphPad Prism 7 (GraphPad Software, San Diego, California), all measurements were done in duplicates.

### Kinetic characterization

The nerve agent substrates were prepared at a small non-hazardous scale, *in situ*, by mixing nontoxic precursors (detailed protocols are available upon request). All kinetic assays were done in 50 mM Tris, 50 mM NaCl, pH 8.0, 25 °C. The rate of hydrolysis of the toxic isomers were determined as described in Ref. [20]. Kinetic parameters for *S*_p_-VX and *S*_p_-RVX were determined by two methods – the loss of AChE activity and the release of thiol group. Thus, rates of hydrolysis of *S*_p_-VX and *S*_p_-RVX were determined by following the release of the thiol leaving group using Ellman’s reagent (DTNB) at 412 nm [26]. Data points were fitted to a mono-exponential function and the apparent first order constant was divided by the enzyme concentration to derive the k_cat_/K_M_ value. The k_cat_/K_M_ values for detoxification of all six nerve agents (*S*_p_-VX, *S*_p_-RVX, *S*_p_-GD, *S*_p_-GF, *S*_p_-GB and *R*_p_-GA) were obtained by monitoring the loss of inhibition of AChE activity in presence of the tested PTE variants. Recombinant human AChE was applied, and its residual activity allowed to quantify the nerve agent concentrations as a function of reaction time. This protocol gave very similar k_cat_/K_M_ values for *S*_p_-VX and *S*_p_-RVX as obtained with DTNB (Table 1). Notably, soman contains two chiral centers with 4 stereoisomers. The two *S*_p_ isomers, namely *S*_P_*S*_C_ and *S*_P_*R*_C_, are toxic and the biphasic detoxification time course (not shown) suggests that their rate of hydrolysis by the PTE cocktail differ by a factor of ~10. However, which isomer is hydrolyzed faster is unknown at this stage since we only used racemic soman as substrate. Rates of hydrolysis of paraoxon were obtained by monitoring the release p-nitrophenol at 405 nm, and k_cat_/K_M_ were derived by fitting initial rates to the Michaelis-Menten model.

**Table 1.**
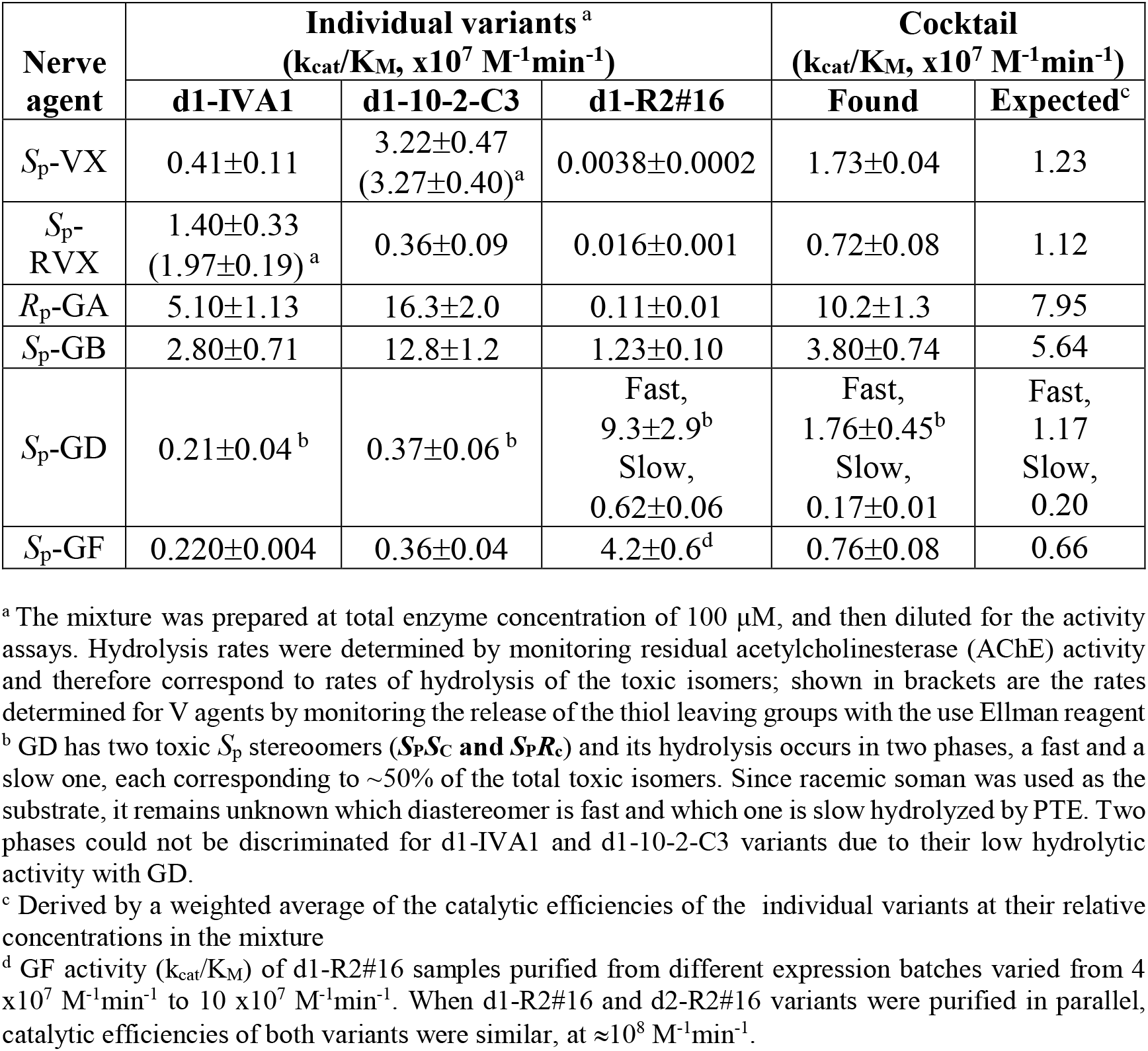
Catalytic efficiency (k_cat_/K_M_, x10^7^ M^−1^min^−1^) of the individual PTE variants and of their mixture for nerve agents detoxification. Shown are k_cat_/K_M_ values for hydrolysis of the toxic isomers, by the three d1-PTE variants, individually, and in a mixture comprising: 60% d1-IVA1, 30% d1-10-2-C3, and 10% d1-R2#16. Measurements were done in duplicates and mean values and standard deviations of the mean are presented.

| Nerve agent | Individual variants<sup>a</sup><br>($k_{cat}/K_M$, $\times 10^7 \text{ M}^{-1}\text{min}^{-1}$) | | | Cocktail<br>($k_{cat}/K_M$, $\times 10^7 \text{ M}^{-1}\text{min}^{-1}$) | |
| --- | --- | --- | --- | --- | --- |
|  | d1-IVA1 | d1-10-2-C3 | d1-R2#16 | Found | Expected<sup>c</sup> |
| $S_p$-VX | 0.41±0.11 | 3.22±0.47<br>(3.27±0.40)<sup>a</sup> | 0.0038±0.0002 | 1.73±0.04 | 1.23 |
| $S_p$-RVX | 1.40±0.33<br>(1.97±0.19)<sup>a</sup> | 0.36±0.09 | 0.016±0.001 | 0.72±0.08 | 1.12 |
| $R_p$-GA | 5.10±1.13 | 16.3±2.0 | 0.11±0.01 | 10.2±1.3 | 7.95 |
| $S_p$-GB | 2.80±0.71 | 12.8±1.2 | 1.23±0.10 | 3.80±0.74 | 5.64 |
| $S_p$-GD | 0.21±0.04<sup>b</sup> | 0.37±0.06<sup>b</sup> | Fast,<br>9.3±2.9<sup>b</sup><br>Slow,<br>0.62±0.06 | Fast,<br>1.76±0.45<sup>b</sup><br>Slow,<br>0.17±0.01 | Fast,<br>1.17<br>Slow,<br>0.20 |
| $S_p$-GF | 0.220±0.004 | 0.36±0.04 | 4.2±0.6<sup>d</sup> | 0.76±0.08 | 0.66 |

<sup>a</sup> The mixture was prepared at total enzyme concentration of 100 $\mu\text{M}$ , and then diluted for the activity assays. Hydrolysis rates were determined by monitoring residual acetylcholinesterase (AChE) activity and therefore correspond to rates of hydrolysis of the toxic isomers; shown in brackets are the rates determined for V agents by monitoring the release of the thiol leaving groups with the use Ellman reagent
<sup>b</sup> GD has two toxic $S_p$ stereoisomers ( $S_pSc$ and $S_pRc$ ) and its hydrolysis occurs in two phases, a fast and a slow one, each corresponding to $\sim 50\%$ of the total toxic isomers. Since racemic soman was used as the substrate, it remains unknown which diastereomer is fast and which one is slow hydrolyzed by PTE. Two phases could not be discriminated for d1-IVA1 and d1-10-2-C3 variants due to their low hydrolytic activity with GD.
<sup>c</sup> Derived by a weighted average of the catalytic efficiencies of the individual variants at their relative concentrations in the mixture
<sup>d</sup> GF activity ( $k_{cat}/K_M$ ) of d1-R2#16 samples purified from different expression batches varied from 4 $\times 10^7 \text{ M}^{-1}\text{min}^{-1}$ to 10 $\times 10^7 \text{ M}^{-1}\text{min}^{-1}$ . When d1-R2#16 and d2-R2#16 variants were purified in parallel, catalytic efficiencies of both variants were similar, at $\approx 10^8 \text{ M}^{-1}\text{min}^{-1}$ .

### Storage stability

Stability of individual variants was measured at two concentrations, according to their concentrations in the mixtures containing 10 μM and 100 μM total PTE active site concentrations. The individual variants, and enzyme mixtures were kept at ambient temperature in storage buffer. Aliquots of the individual stored samples were diluted to 1 nM in 50 mM Tris pH 8.0 and 100 mM NaCl, and the activity with 0.18 mM paraoxon was monitored. Initial rates were derived and compared to the initial rate at the onset of incubation (time zero). Assays of the cocktail stability at ambient temperature were performed by dilution of stored 10 μM and 100 μM stocks to 1 µM total enzyme concentrations. The final PTEs concentration in the assay reactions were 25 nM for VX and RVX (DTNB protocol) and 40-50 nM for GF (detoxification using AChE). The k_cat_/K_M_ of the incubated enzymes were calculated as above.

### pH-activity assay

Variants (0.4 µM) were incubated in 100 mM sodium acetate buffer (pH 5.0 – 6.0), 100 mM NaCl and 0.1 mM ZnCl_2_, for 1.5 h at the ambient temperature. The incubated samples were first diluted 100-fold in storage buffer, and additionally diluted 10-fold in the activity buffer with 0.18 mM paraoxon. The initial rates were derived as above and normalized to the rate of an enzyme sample stored in activity buffer (pH 8.0).

## Results

### Engineering tag-less variants

The three PTE variants applied here, d1-IVA1, d1-10-2-C3 and d2-R2#16, were stabilized using computational design algorithm-PROSS [27]. Nonetheless, these variants were all expressed as a fusion protein with N-terminal maltose-binding protein (MBP). Fusion to MBP prompted higher expression yields as well as more reproducible expression levels in *E. coli* cells grown in 96-well plates, and was thus helpful for library screens. However, the MBP fusion has several major drawbacks with respect to application in decontamination or/and detoxification. Purification based on affinity chromatography of MBP is costly, and over half of the produced protein mass is redundant. *In vivo* administration of yet another protein (*i.e.*, MBP in addition to PTE) is problematic. Foremost, we observed that upon storage of MBP-PTE fusions, spontaneous cleavage of the connecting linker occurs thus yielding a heterogeneous protein mixture (Figure S2). A cleavable MBP tag is an option, but this production protocol is laborious and costly. We also observed that to separate the tag-free PTE cleavage must be near-complete because PTE is a dimer, and thus the tag of both subunits must be cleaved. We therefore sought to express and purify these PTE variants as intact, *i.e.*, not only without a fusion protein, but also without any tag, thus reducing production cost and protein antigenicity.

We therefore cloned and expressed wild-type PTE, and the three engineered variants without a signal sequence and with no other additions at either the N- or the C-termini (amino acids and DNA sequences are provided in Supplementary Table S1). Whereas, in this format, wild-type PTE gave very low expression, the engineered variants, likely owing to the PROSS optimization, expressed at ≥ 40-fold higher levels (the latter also had optimized codon sequences). A simple purification protocol was developed based on ion-exchange chromatography, allowing us to obtain the three engineered variants at relatively high yield and purity (≥ 95 % as judged by SDS-PAGE, Figure S1).

Comparison of the kinetic parameters indicated that the intact variants exhibited comparable, if not better, catalytic efficiency compared to their MBP-fusions (Table S2). Their thermostability, as judged by heat inactivation assays was above 60 °C (Figure 2A) and similar to the thermal stability of the corresponding MBP variants. However, the heat inactivation curves of the MBP fusions did not fit a single sigmoidal curve, but were rather biphasic (Figure 2B). One sigmoid largely overlapped the untagged variant while the second one indicated a protein species with lower thermostability. This biphasic heat inactivation curve seems to be the outcome of the heterogeneity of the MPB-PTE protein preparations. Heterogeneity was indeed observed in these preps, probably due to instability of MBP-tag (*i.e.*, due to a mixture of PTE with and without tag; Figure S2) and/or due to coexistence of protamers and dimers (discussed below). The biphasic heat inactivation curves suggest that MBP-fused and cleaved PTE have different melting temperatures. The former exhibits lower stability, possibly because unfolding of the MBP tag may induce earlier unfolding of the fused PTE.

**Figure 2.**
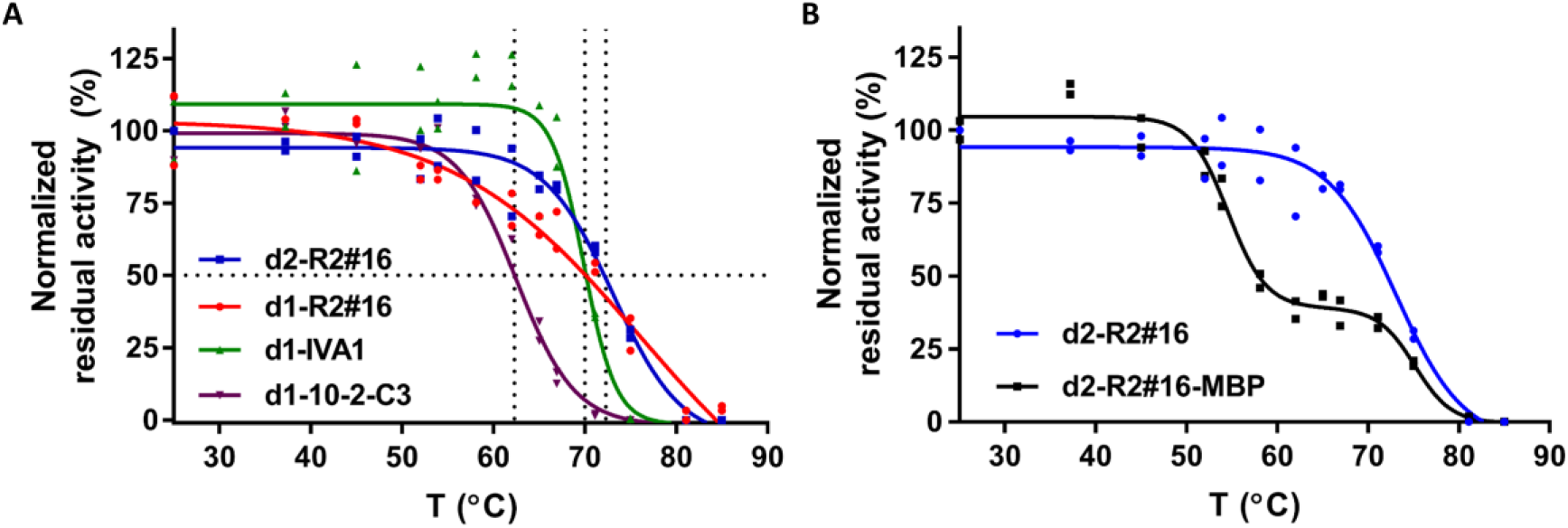
Thermostability of the engineered PTE variants. **A.** Heat inactivation assay of the tag-less variants. The variants (at 2.8 µM; concentrations are given per PTEs active sites) were incubated at different temperatures for 30 minutes, and their residual activity was tested with paraoxon and normalized to the activity of the same variant incubated at 25 °C. **B**. Heat inactivation of d2-R2#16 with and without an MBP-tag. Measurements were done in duplicates and values normalized to the average activity at 25 °C are presented.

### Engineering higher sequence identity

We performed another engineering step toward the desired enzyme mixture. As it turned out, variants d1-IVA1 and d1-10-2-C3 have the same PROSS background with 9 stabilizing mutations (d1-PTE), while variant d2-R2#16 has a different PROSS background with 19 stabilizing mutations (d2-PTE; see Table 3 in Ref. [27]). To increase the homogeneity of this enzyme mixture, and also enable potential application of this enzyme cocktail as the single drug, we changed PROSS background of d2-R2#16 variant to d1-PTE. In doing so we increased sequence identity between the three variants from 92% to above 96 %.

Crucially, the catalytic efficiency with respect to GF, the nerve agent that comprises R2#16’s primary target, did not change. The melting temperature of the new d1-R2#16 variant decreased by nearly 10 °C as judged by fluorescence measurements (Figure S3). However, its thermal stability as measured by loss of activity was similar to d2-R2#16 (Figure 2A). Overall, despite carrying many active-site mutations, the melting temperature of all the engineered tag-less variants was above 60 °C. An additional potential advantage of d1-R2#16 is its resistance to acidic pH that is higher compared to d2-R2#16 although both these variants are less acid-tolerant than d1-10-2-C3 (pH< 5.3; Figure S4). Resistance to low pH provides improved functionality at high nerve agent concentrations (OPNA hydrolysis releases acid, and decontamination is typically local, *e.g.* drops of neat agents, and buffering has therefore limited capacity).

### Nerve agent degradation

The intact PTE variants, d1-10-2-C3, d1-IVA1 and d1-R2#16, were individually produced and purified. Based on their individual performances, a mix of 60:30:10 molar ratio of d1-IVA1, d1-10-2-C3 and d1-R2#16, respectively, should, in theory, enable efficient hydrolysis of all six threat nerve agents with k_cat_/K_M_ ~ 10^7^ M^−1^ min^−1^. The rate of degradation of the toxic stereoisomers were determined by monitoring the loss of acetylcholinesterase inhibition, for both the three individual variants and for their mixture. The rate of V-agents hydrolysis was also determined by monitoring the thiol leaving group appearance using the Ellman reagent. This assay monitors hydrolysis of both stereoisomers, however, we employed the stereo-specifically synthesized *S*_p_ enantiomers of the V agents.

Overall, these assays indicated that the 3-variants mixture exhibits effective detoxification of all six threat agents (Table 1). The rates matched the rates expected from the rates of the individual variants, thus indicating that mixing caused no interference. The mixture’s catalytic efficiency of detoxification, namely k_cat_/K_M_ with the enzyme concentration taken as the total concentration of the three enzymes, was in the range of 10^7^ M^−1^min^−1^ for all tested agents, except for one of the soman isomers that exhibited a k_cat_/K_M_ of 0.17×10^7^ M^−1^min^−1^. However, different mixtures could obviously be made in which d1-R2#16 is added in proportions higher than 10%, thus allowing better protection against soman.

### Stability of the 3-variants mixture

One specific concern regards the fact that PTE is a dimer. In wild-type, the dimer is relatively stable, although at low concentration (< 1 nM), and in the absence of a stabilizing protein such as bovine serum albumin, activity is rapidly lost. In some engineered variants, especially those with mutations in loop 8 (residues 307 – 312) that comprises part of the dimer interface, the dimer may be further compromised. In a mixture of variants, dimer dissociation would also lead to hetero-dimers that might exhibit low activity.

To test for loss of activity upon long storage, due to possible hetero-dimers formation, or any other reason, we measured in parallel the activity of the individual variants, and their corresponding mixture at the same concentration, upon storage at room temperature for 4 weeks (Figure 3). The individual variants exhibited no, or only minor loss of activity throughout the four weeks (Figure 3A). The most significant loss of activity occurred with respect to GF in the variant mixture (Figure 3B). Notably, the GF activity decreased ~2-fold several hours after mixing, and then remained intact in the next 4 weeks, suggesting that d1-R2#16 the variant which confers the overwhelming majority of the GF hydrolyzing activity is affected by mixing. The loss of ~50% of the GF hydrolyzing activity was reproducibly observed, also when concentrations were increased 10-fold (total enzyme concentration of 100 µM in the cocktail mixture). Overall, it appears that while mixing these three variants causes a partial loss of activity, this loss is tolerable, and can obviously be readily compensated by higher enzyme dose.

**Figure 3.**
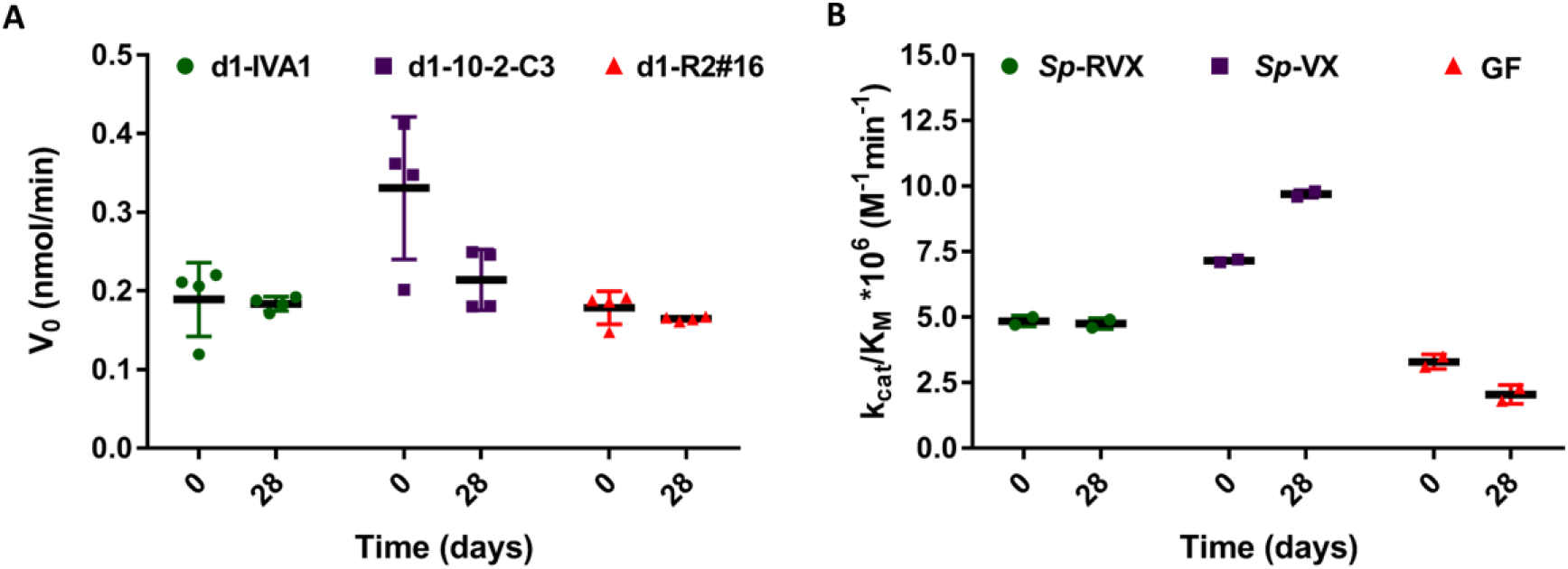
Stability of the individual variants and of their mixture to long-term storage at ambient temperature. **A.** The residual activity of the individual variants was measured with paraoxon following 28 days of storage at ambient temperature (22-25 °C). Storage concentration of the individual variants was identical to their concentration in the 10 µM mixture: 6 µM d1-IVA1, 3 µM d1-10-2-C3 and 1 µM d1-R2#16. For monitoring paraoxon activity variants were diluted to 1 nM in the reaction buffer (shown are intial rates measured in this activity assay). **B**. Activity of the variant mixture, at a total enzyme concentration of 10 µM, containing 6:3:1 ratio of d1-IVA1, d1-10-2-C3 and d1-R2#16, respectively, after 28 days of incubation at ambient temperature. Cocktail activity was monitored with 3 different substrates, *S*_p_-RVX (reporting mostly d1-IVA1 activity), *S*_p_-VX (reporting mostly d1-10-2-C3 activity) and racemic GF (reporting d1-R2#16 activity towards the toxic *S*_p_-GF). The collected data points are shown, with their mean values depicted as horizontal lines and standard deviation as vertical lines (panel A, n=4, and panel B, n=2).

## Concluding remarks

We describe a cocktail of three PTE variants with 96% identical sequences that efficiently detoxifies the entire range of known threat nerve agents. We have also optimized expression and purification of the cocktail variants to afford stable proteins at relatively low cost. These PTE variants are highly stable, also as a mixture, even when stored in solution for one month at ambient temperature. To our knowledge, this engineered ‘cocktail’ comprises the first solution for effective enzymatic detoxification and/or decontamination of the entire range of known threat nerve agents, including both G- and V-type agents.

## Abbreviations

OP: organophosphate
PTE: phosphotriesterase
OPNA: organophosphate nerve agent
OPH: organophosphate hydrolase
RVX: Russian VX

## Acknowledgments

We gratefully acknowledge financial support from Defense Threat Reduction Agency (DTRA) of the US Department of Defense (HDTRA1-17-0057). D.S.T. is the Nella and Leon Benoziyo Professor of Biochemistry. We are grateful to Kesava Phaneendra Cherukuri for assistance in making Figure 1.

## Supplementary Information

**Figure S1:**
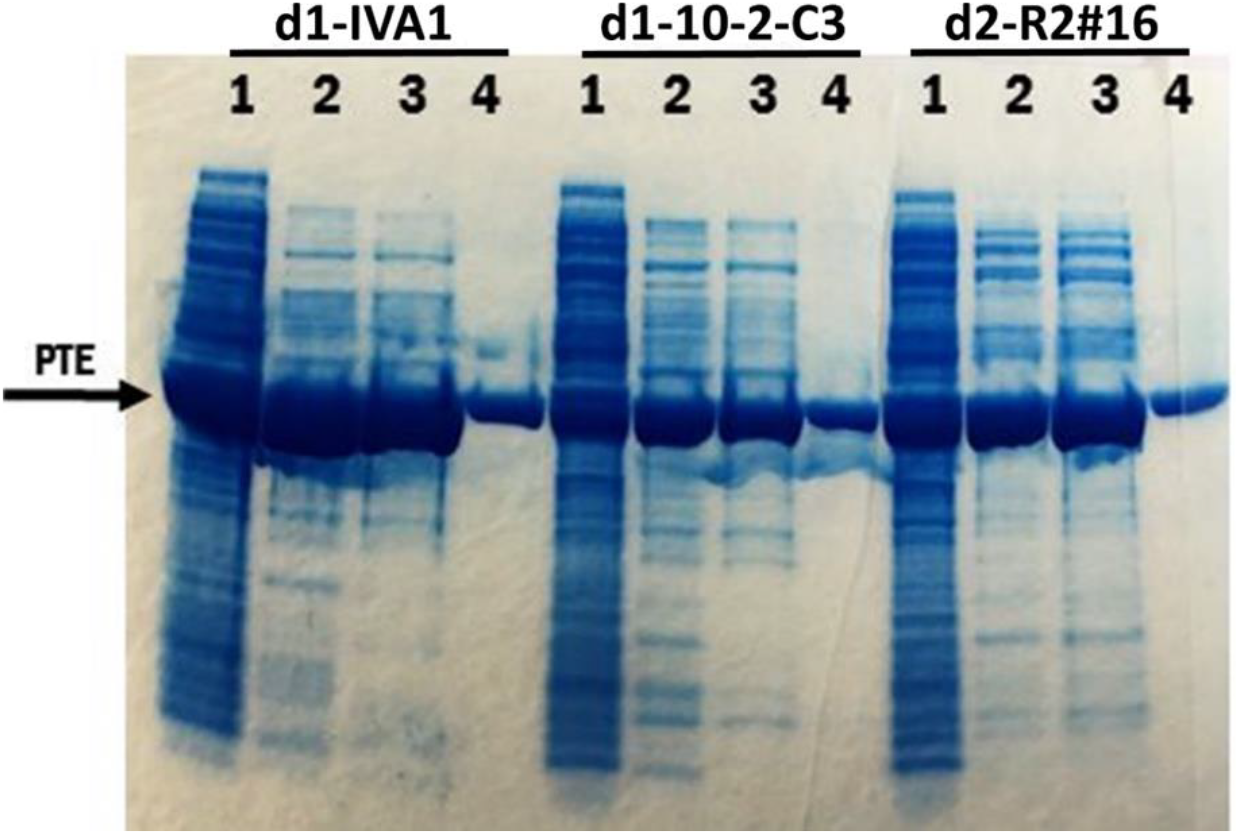
Three-step purification of the tag-less PTE variants. #1-cell lysate; #2 - after anion chromatography (1^st^ purification step), #3 - after cation chromatography (2^nd^ step), #4 - after hydrophobic interaction chromatography (3^rd^ step).

**Figure S2:**
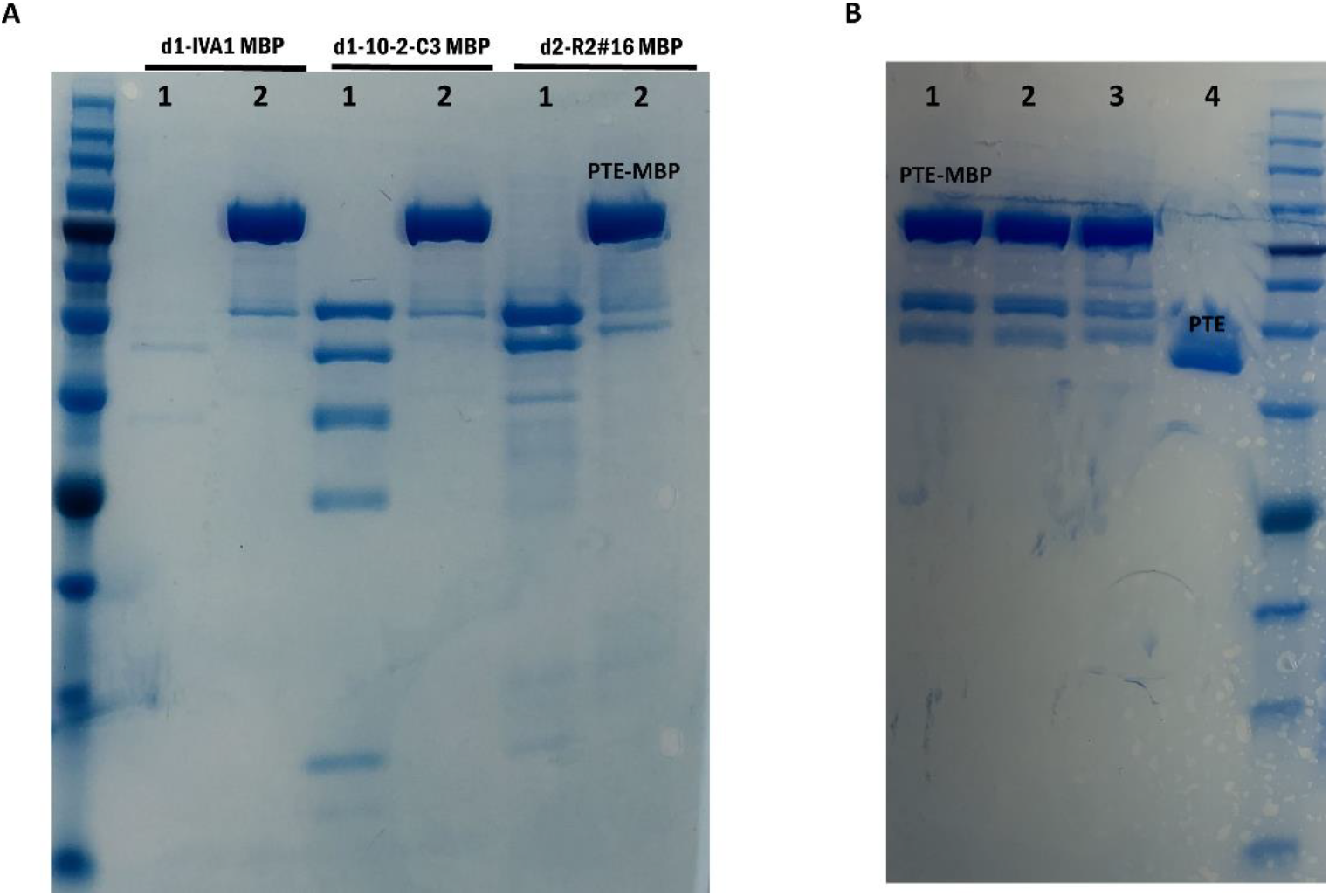
PTE-MBP degradation during storage. **A.** Comparison of PTE variants immediately after purification (Lanes #2) and after long storage (Lanes #1;d1-IVA1 MBP and d1-10-2-C3 MBP after 3 years, and d2-R2#16 after 6 months). The concentration of the variants was set to 10 µM except 3 years old d1-IVA1 MBP sample (0.44 µM). **B.** MBP-PTE variants after 1 year of storage at 4 °C (the same freshly purified variants are in Lanes #2 gel in panel A); 1 - d1-IVA1, 2 - d2-10-2-C3, 3 - d2- R2#16 and 4 – tag-free d1-IVA1 variant. The concentration of the variants was set to 10 µM.

**Figure S3.**
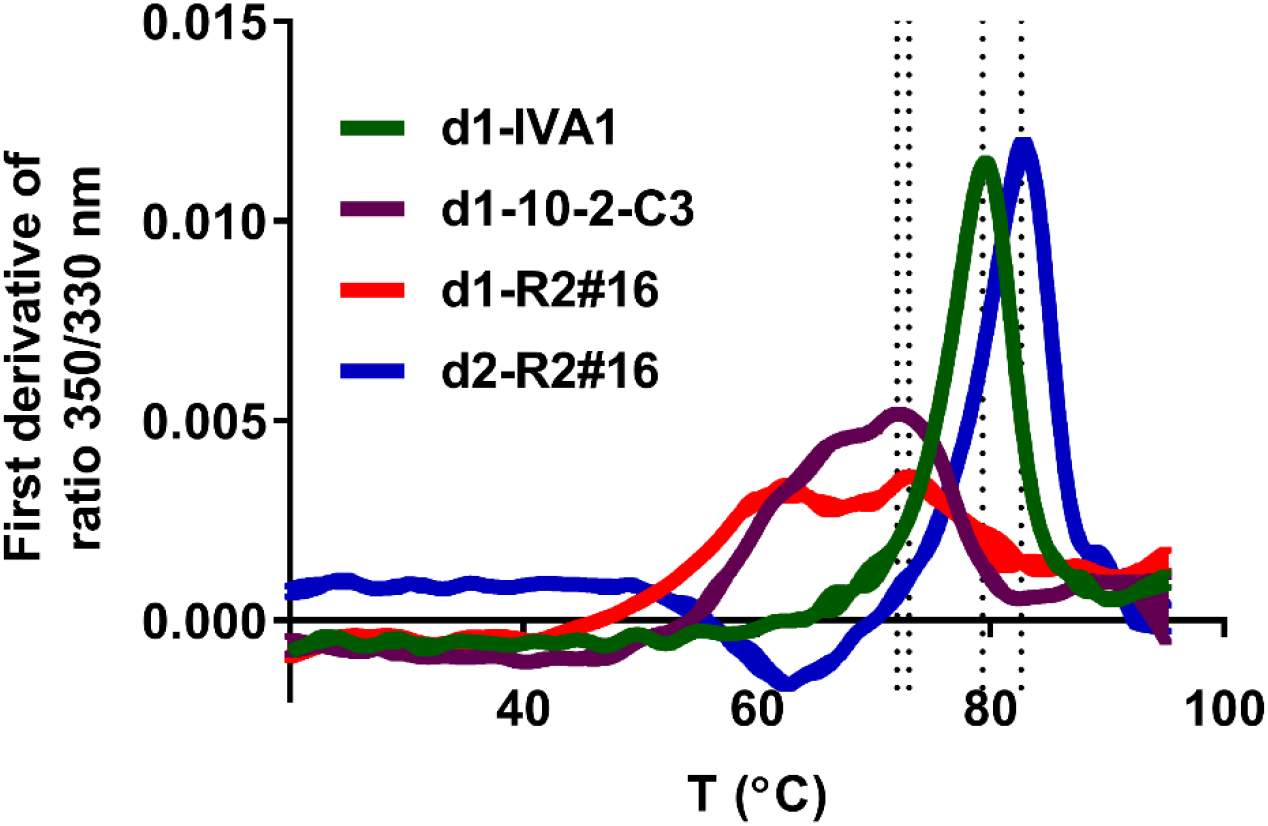
Melting curves of the engineered PTE variants. Protein unfolding was monitored by the change of fluorescence intensity of tryptophan or tyrosine (excitation at 280 nm, emission at 350 and 330 nm) at increasing temperatures using Prometheus NT.48, Nano Temper. Protein concentrations were 8.3 µM. T_M_, the temperature at which 50 % of the sample appears to be unfolded (dotted lines), was derived from the instrument’s fit: d1-IVA1, 79.3 °C; d1-10-2-C3, 72.0 °C; d1-R2#16, 73.0 °C; d2-R2#16, 82.6 °C.

**Figure S4.**
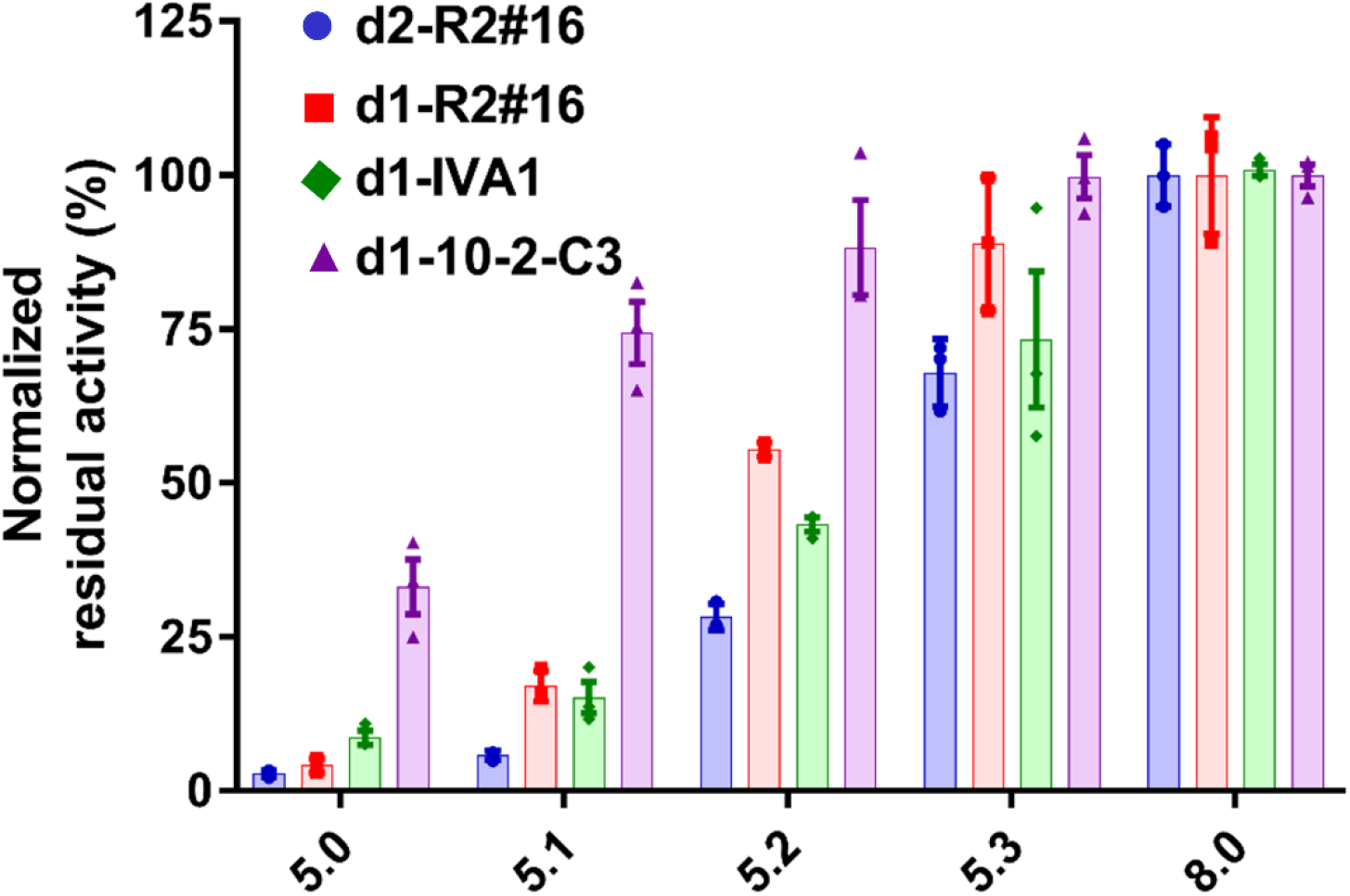
pH-activity profile of the engineered PTE variants. Variants (0.4 µM) were incubated for 1.5 h at various pH’s and their residual activity was measured with paraoxon in activity buffer pH 8.0. Residual activity was normalized to the activity of protein sample incubated in storage buffer at pH 8.0. Measurements were done in triplicates and the collected data points are shown, with their mean values depicted as rectangular and standard error of the mean as vertical lines.

**Table S1:**
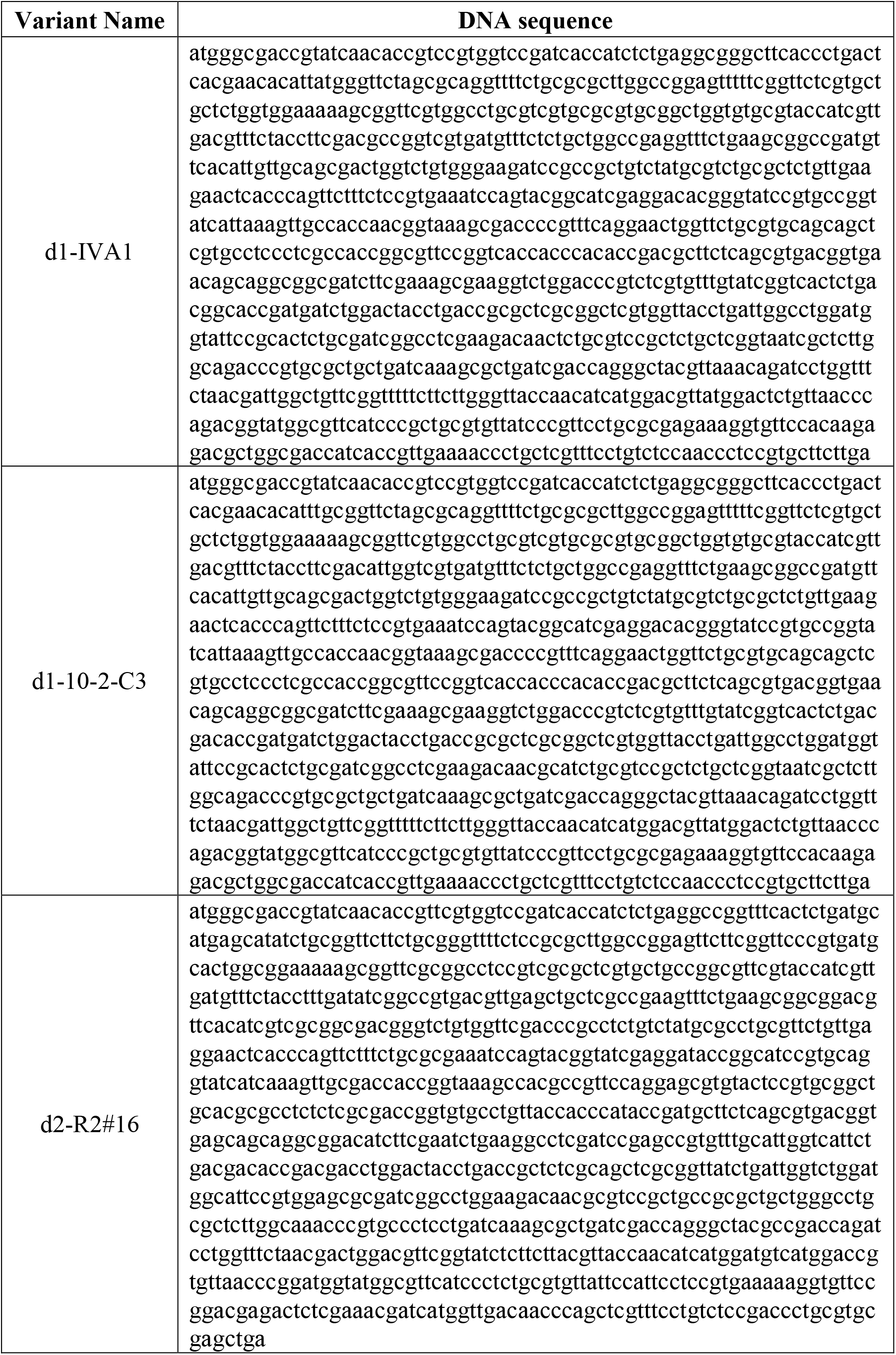
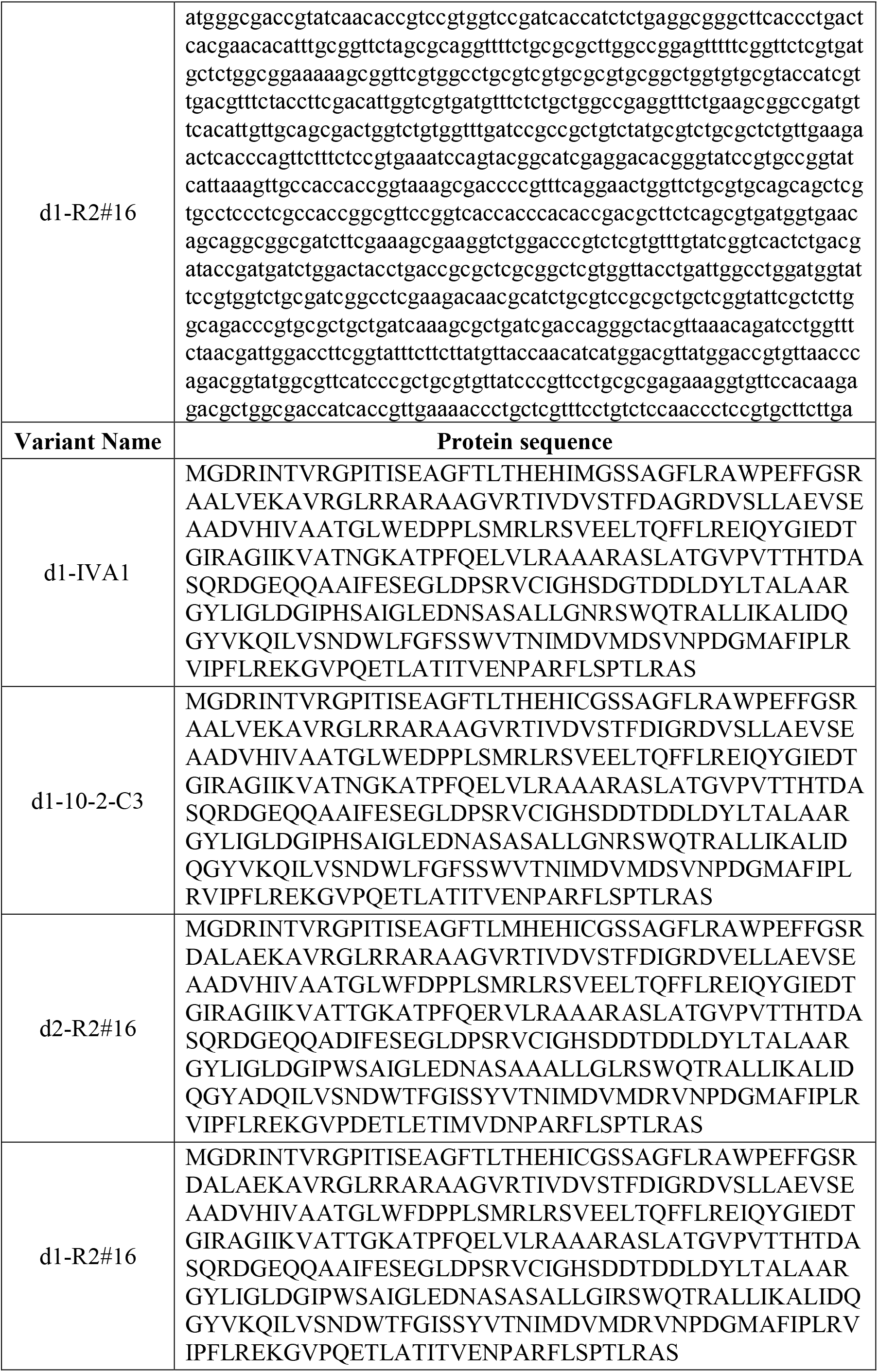
DNA and protein sequences of the PTE variants described here.

**Table S2.**
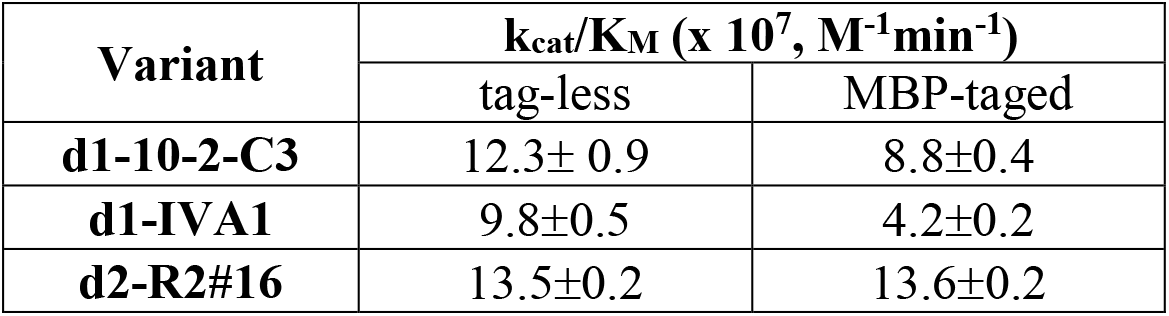
k_cat_/K_M_ values (x 10^7^, M^−1^min^−1^) of the engineered PTE variants with paraoxon. Measurements were done in duplicates and mean values with standard of the mean are presented.

